# Declining reservoir elevations following a two-decade drought increase water temperatures and non-native fish passage facilitating a downstream invasion

**DOI:** 10.1101/2024.01.23.576966

**Authors:** D.E. Eppehimer, C.B. Yackulic, L.A. Bruckerhoff, J. Wang, K.L. Young, K.R. Bestgen, B.A. Mihalevich, J.C. Schmidt

## Abstract

River ecosystems are threatened by interactions among river regulation, non-native species, and climate change. Water use has exceeded supply for two decades in the USA’s Colorado River basin draining its two largest storage reservoirs (Lake Powell and Lake Mead). In 2022, after more than two decades of declining reservoir levels and warming downstream river water temperatures, Lake Powell began releasing water from its lower epilimnion into the Grand Canyon segment of the Colorado River. While managers were concerned about the risk of non-native, predatory smallmouth bass (*Micropterus dolomieu*) entrainment and reproduction, they lacked a quantitative tool to understand entrainment rates and population growth potential under different reservoir management strategies. To fill this void, we developed models in spring 2022 that: 1) predicted propagule pressure at different reservoir elevations, and 2) linked reservoir storage/operations, water temperatures, and smallmouth bass population dynamics to forecast population growth rates under different hydrologic and management scenarios. In the summers of 2022 and 2023, smallmouth bass were documented reproducing in the lower Colorado River for the first time. Our models accurately forecasted adult catch of smallmouth bass in 2022 and 2023 and forecasted that reproduction would occur in both years for the first time in the history of this river segment. Above average runoff in 2023 increased reservoir elevations, however the potential for smallmouth bass establishment remains high because of long-term forecasts of reduced reservoir inflows and lake levels significantly below full pool. Maintaining Lake Powell elevations above 1,088 m (3,570 ft) would likely minimize propagule pressure from the reservoir and would likely create downstream conditions that minimize smallmouth bass population growth.

## Introduction

The spread of non-native species can lead to biodiversity losses and economic losses, and prevention or early intervention is generally considered the most effective approach to manage the spread of non-native species (Pimentel et al., 2005). Non-natives species can alter ecosystems and may drive declines in native species diversity and abundance (Simberloff et al., 2013; Gallardo et al., 2016, Blackburn et al., 2019). The economic cost of resource damage from and management cost of non-native invasions is high (Cuthbert et al., 2021). For example, the USA is estimated to have spent $23 billion controlling/managing aquatic invasive species in 2020 alone (Cuthbert et al., 2021). Management success varies dramatically, but invasion prevention and/or early responses are most effective and are typically cheaper than long-term management of established invaders (Pimentel et al., 2005; Larson et al., 2011). Predictive models are helpful for evaluating potential management strategies as preventative or early management responses are frequently absent or inadequate when invasion forecasts are unavailable (Runge et al., 2018; Healy et al., 2023).

Smallmouth bass (*Micropterus dolomieu*) are a popular sport fish that have been stocked around the globe but can negatively impact native freshwater ecosystems (reviewed in Loppnow et al., 2013). Smallmouth bass are particularly successful invaders due to their tolerance to a range of environmental conditions, early piscivory, and relative high survival of eggs/larvae due to parental care (Brown et al., 2009). Smallmouth bass are capable of inhabiting both lakes and streams, and once introduced they often spread from introduction points, which are typically in or below reservoirs, and their expansion in regulated rivers is facilitated by altered flow, thermal, and sediment regimes (Havel et al., 2005). Smallmouth bass are identified as a threat to native fishes of Western USA river basins (Johnson et al., 2008), and have spread across the Colorado River basin with their increases in abundance often coinciding with declines in federally listed native fishes (Dibble et al., 2021). Concerns that smallmouth bass populations have had negative impacts on native fish in the upper Colorado River basin (upstream of Lake Powell) have motivated active smallmouth bass removal programs for more than ten years. These efforts have shown temporary reductions in populations but have failed to suppress smallmouth bass over longer time periods when environmental conditions are suitable for reproduction (Breton et al., 2014) and has not led to increases in federally listed fish populations that provide the motivation for removals.

The ongoing Millennium Drought and water supply crisis in the Colorado River basin (Schmidt et al., 2023) threatens water security and native ecosystems (Schmidt et al., 2022). Water supply and storage decisions change river suitability for fishes including non-native species (Dibble et al., 2021; Bruckerhoff et al., 2022), and recent, low reservoir elevations have facilitated the spread of non-native species like smallmouth bass. Reservoir elevations change the temperature of dam releases in systems with fixed intake elevations, potentially altering thermal suitability for fishes downstream (Dibble et al., 2021). Reservoir elevations also change rates of fish passage through dams as fish are not distributed uniformly with respect to depth in reservoirs (Harrison et al., 2019). Smallmouth bass have been present in the upper Colorado River basin since the late 1960s through reservoir stockings. Lake Powell, the reservoir on the Colorado River upstream from Grand Canyon, formed by Glen Canyon Dam (Fig. 1A), was stocked with smallmouth bass from 1982 to 1989 (Pennock and Gido, 2021); the initial stocking was unauthorized (Goldfarb, 2023). Prior to 2022, smallmouth bass passage through Glen Canyon Dam was likely rare as smallmouth bass are concentrated in the upper ∼14 m of the reservoir and the fixed intakes were much deeper below the reservoir surface (see Supplemental Materials). Furthermore, release temperatures from Glen Canyon Dam were likely too cold for reproduction by adults that were entrained (Dibble et al., 2021). However, in 2022 after more than ten years of declines, Lake Powell reached its lowest elevation in over 50 years (Fig. 1B), and epilimnetic water releases were >4°C warmer than the maximum temperature observed in those prior 50 years (USGS Stream Gage: 09380000; USGS, 2023). For multiple months, water temperatures exceeded 16°C, which is the estimated spawning threshold for smallmouth bass (see Supplemental Materials; Table S1). Subsequently, smallmouth bass reproduction was observed below Glen Canyon Dam, which has never been previously reported in this river segment. This smallmouth bass invasion threatens native fishes, including federally listed humpback chub (*Gila cypha*), as well as the tailwater rainbow trout (*Oncorhynchus mykiss*) fishery. Over the last 15 years, humpback chub in the Grand Canyon have increased dramatically in abundance (Yackulic et al., 2014; Van Haverbeke et al., 2017; Dzul et al., 2023) leading to their downlisting from US Endangered Species Act “endangered” classification to “threatened” in 2021 (USFWS, 2021). However, in the early 2000s, one of six recognized humpback chub populations was functionally extirpated coincident with increases in smallmouth bass and another federally listed fish species had continued to decline coincident with smallmouth bass populations increases (Dibble et al., 2021). Managers are concerned whether a smallmouth bass invasion of the Grand Canyon might lead to similar declines in humpback chub, therefore reversing recent species recovery.

**Figure 1:**
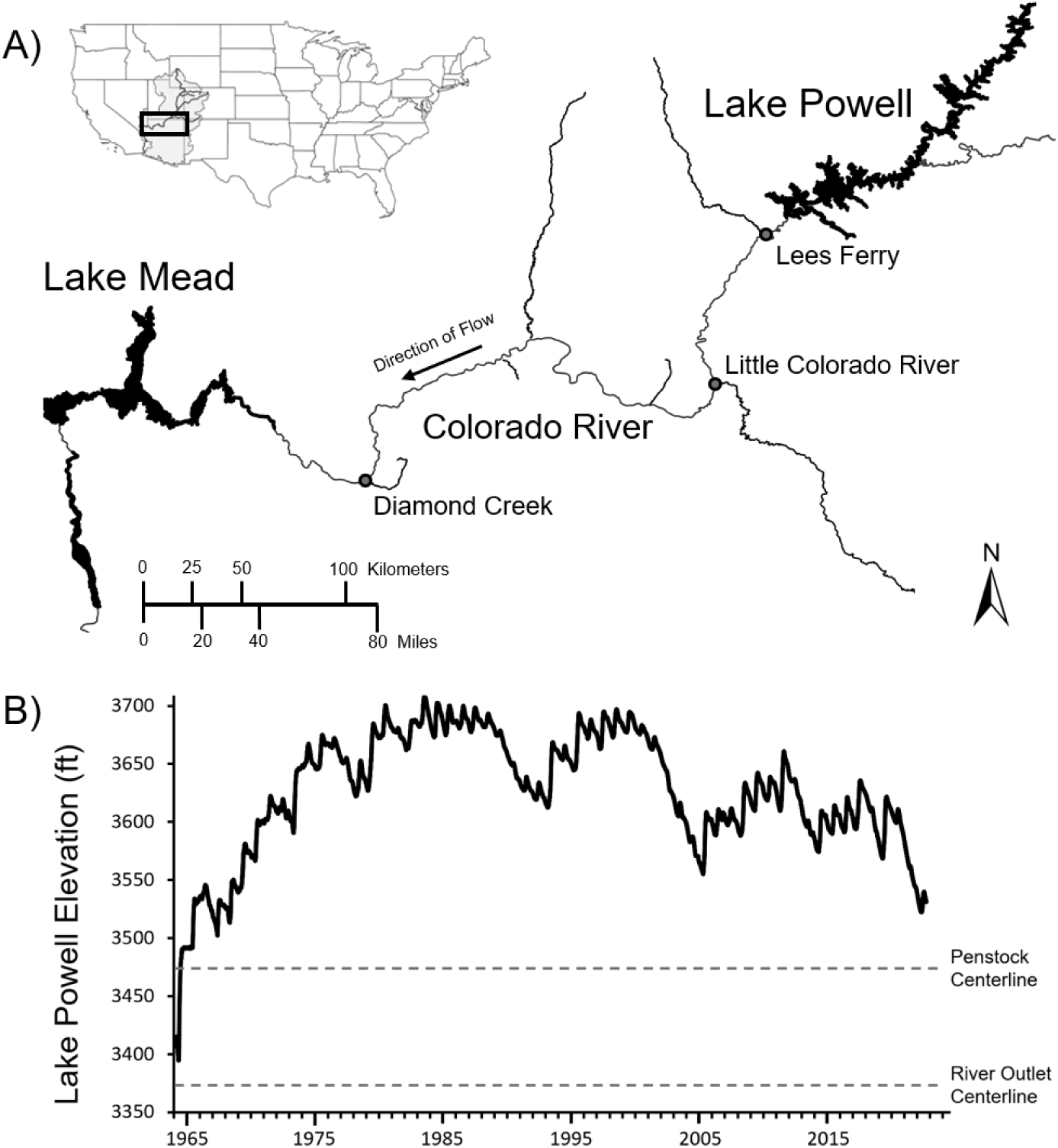
Study site in the context of historical variation in Lake Powell elevation. A) Map of the Colorado River between Lake Powell and Lake Mead. The Lees Ferry, Little Colorado River confluence, and Diamond Creek confluence locations are denoted with grey circles. B) Historic Lake Powell elevation above sea level in feet since dam closure. Grey dashed line denotes the centerline elevation of Glen Canyon Dam penstock and river outlet intakes.

To inform ongoing management decisions, we developed models to forecast smallmouth bass invasion risk under different reservoir storage and dam operations. First, we developed a model that estimates smallmouth bass emigration from Lake Powell under different reservoir elevations to identify thresholds for increased fish entrainment and passage through Glen Canyon Dam. Second, we coupled reservoir operations with water temperature and smallmouth bass population models to predict smallmouth bass population growth rate (λ). These models were developed in spring of 2022 (prior to observations of reproduction), and we evaluate model performance based on observed catch of adults via regular monitoring and whether reproduction was observed in 2022 and 2023. We also illustrate the utility of these models by forecasting entrainment and population growth rate over a range of potential future conditions to inform ongoing management decisions regarding reservoir operations.

## Methods

### Study site

Our study is focused on the segment of the Colorado River flowing through Glen, Marble, and Grand Canyons in Arizona, USA, and bounded by two reservoirs (Lake Powell and Lake Mead). Water from Lake Powell is released through Glen Canyon Dam and travels approximately 475 river kilometers (rkm) before entering Lake Mead with environmental conditions and aquatic communities changing dramatically over its length. Lees Ferry, the tailwater section located in the first 25 rkm downstream of Glen Canyon Dam, is characterized by clear water, abundant aquatic vegetation, and water temperatures almost entirely dependent on reservoir release temperatures (Mihalevich et al., 2020). Since 1973, this reach was characterized by three decades of very cold water temperatures (95% range of daily water temperatures: 7.6 °C – 11.3 °C), followed by approximately two decades of cool water temperatures prior to 2022 (95% range of daily water temperatures: 8.1 °C – 14.3 °C) (USGS stream gage: 09380000; USGS, 2023) and was managed as a blue-ribbon rainbow trout fishery (Runge et al., 2018). Below Lees Ferry, the river becomes more turbid, aquatic vegetation becomes rare, and water temperatures gradually become less impacted by Glen Canyon Dam. Approximately 122 rkm below the dam, the Colorado River reaches its confluence with the Little Colorado River, a tributary serving as a population center for humpback chub in Grand Canyon (Yackulic et al., 2014). Further downstream in western Grand Canyon, near the confluence of the Colorado River and Diamond Creek (∼386 rkm downstream from Glen Canyon Dam), humpback chub populations have increased dramatically over the last decade (Van Haverbeke et al., 2017; Dzul et al., 2023). Our models characterize smallmouth bass propagule pressure from Lake Powell to Lees Ferry, as well as the potential for smallmouth bass population growth based on Colorado River temperatures at Lees Ferry, the Little Colorado River confluence, and Diamond Creek confluence (Fig. 1A).

### Glen Canyon Dam operations and reservoir elevations

Glen Canyon Dam can release water through penstocks and river outlets (Fig. 1B). Typically, water is passed through the penstocks at a centerline elevation of 3,470 ft (1,058 m) to generate hydropower; turbines likely reduce survival rates but do not prevent fish passage (see Supplemental Materials). Water can also be released through the deeper river outlets (centerline of 3,370 ft: 1,027 m). However, the river outlets do not generate hydropower, so their use is rare. Results presented here assume all water is passed through the penstocks until reservoir elevations are at or below 3,490 ft (1,064 m), at which point releases switch to river outlets to avoid damaging hydropower infrastructure, and the reservoir is below ‘power pool’.

Both the propagule pressure and population growth models rely on Lake Powell elevations which we estimated using a reservoir elevation model used by Colorado River basin managers (Schuster, 1998). Changes in Lake Powell elevation were modelled monthly as a function of starting elevation, inflows, outflows, bank storage, and evaporative losses (see Supplemental Materials). For our analyses, we considered a range of future conditions using combinations of starting elevations, inflows, and outflows. Starting elevations ranged from 3,470 (1,058 m and 12% of capacity: below power pool) to 3,600 ft (1,097 m and 47% of capacity). Inflows were based on resampling the past 23 years (2000-2022; Eppehimer et a., 2024). Inflows during this period varied from 4 to 15 million-acre feet per annum (4,934-18,502 million cubic meters per year). In our analyses we assumed one of two scenarios for outflows: 1) 7.48 million acre feet (maf: 8,634 million cubic meters) per year outflows with monthly patterns derived from the Bureau of Reclamation’s 2024 projections (see Supplemental Materials) and 2) a potential management scenario in which reservoir levels are held constant over the course of a year by matching monthly dam release volumes with monthly inflow volumes (this scenario will be referred to as “maintain elevation”). For some analyses, we also produced forecasts for 2024 using the 2024 calendar year projected Lake Powell starting elevation of 3,565 ft (1,087 m), 2024 predicted outflows, and 23 historic inflow traces (2000-2022) characterizing inflow variability.

### Entrainment model fitting and forecasting

We modeled propagule pressure (defined as the number of individuals that are entrained and survive to a particular time point) based on estimates of smallmouth bass densities in the Glen Canyon Dam forebay of Lake Powell, movement, use of different depths, survival during entrainment, adult survival, estimated capture probabilities for Lees Ferry, and annual catch of likely entrained smallmouth bass from 2010 to 2020 (see Supplemental Materials). We fit the model to available data using a Bayesian approach implemented in program Stan version 2.3 (Stan Development Team, 2022) run in R version 4.2.1 (R Core Team, 2021), incorporating literature derived priors for parameters when possible. We then predicted the number of entrained smallmouth bass under different time series of reservoir elevations over the course of 1 and 2-year periods based on different elevations (see Supplemental Materials for details). As part of our model evaluation, we compared the predicted change in catch of adult (>200 mm) smallmouth bass in 2022 and 2023 to actual catches from regular monitoring (unpublished data: L. Tennant, NPS). We focused on adult smallmouth bass because, all but two fish captured before 2022 in Lees Ferry were > 200 mm, and following observed reproduction in 2022, the provenance of fish < 200 mm was unknown (length frequency histograms suggested these locally produced fish did not reach adult sizes in 2022 or 2023).

### Forecasting population growth rate (λ) based on thermal regimes

To predict potential smallmouth bass population growth under different reservoir operations, we used Lake Powell data from 2000 through 2022 (Andrews and Deemer, 2022) to estimate daily water temperatures at one-ft (0.3-m) resolution near the Glen Canyon Dam forebay (see Supplemental Materials). Lake Powell reservoir dynamics are advection driven by each year’s spring inflows (Fig. S1), so our model used both depth and inflow volume to predict water temperature profiles. We interpolated monthly reservoir elevations to daily reservoir elevations and predicted release temperature based on the water temperature in the associated thermal profile at the depth of the intake (either penstocks or river outlets). Water temperature at locations downriver from the dam was then estimated using an existing model adapted to the daily scale (Dibble et al., 2021; see Supplemental Materials).

Our population growth model assumes temperature is the only factor influencing smallmouth population growth (e.g., the model assumes there is no spawner limitation and that there are no habitat or prey availability limitations), and location-specific lambdas (λ) are estimated by calculating the dominant eigenvalue associated with a matrix population model in which production of age-0 fish is temperature dependent with all other vital rates held constant using estimates from portions of the upper Colorado River basin (Breton et al., 2015; unpublished data: L. Bruckerhoff, OSU; see Supplemental Materials). We assumed a spawning initiation temperature lower threshold of 16°C (see Supplemental Materials; Table S1). In the model, temperature directly impacts the occurrence and timing of spawning, the growth of age-0 fish, and subsequent indirect survival of these age-0 fish. All population growth forecasts were run in R (version 4.2.1; R Core Team 2021).

## Results

Smallmouth bass propagule pressure is predicted to increase with decreasing reservoir elevation until 3,490 ft (1,064 m), after which the dam switches to deeper river outlets lowering propagule pressure (Figs. 2; S2). At a Lake Powell elevation of 3,540 ft (1,079 m), the number of propagules is nearly double that at 3,600 ft (1,097 m), however the absolute number is still relatively small. The number of propagules increases dramatically as reservoir elevation declines below 3,540 ft (1,079 m). Decisions regarding the volume of annual releases can have profound impacts on rates of entrainment under certain starting elevations and inflow volumes (Fig. 2). Reservoir elevations in 2022 were the lowest in over 50 years averaging 3,530 ft (1,076 m) and reaching a minimum of 3,522 ft (1,074 m) (Fig. 1B). Reservoir elevations in 2023 reached a lower minimum of 3,520 ft (1,073 m) but increased dramatically due to large snowmelt runoff and average 3,555 ft (1,084 m) over the calendar year. The entrainment model predicted a modest increase in the abundance of adults in Lees Ferry in 2022 coincident with lowered elevations and steady abundances in 2023 and observed catch from regular monitoring in 2022 and 2023 was consistent with model forecasts (Fig. 3).

**Figure 2:**
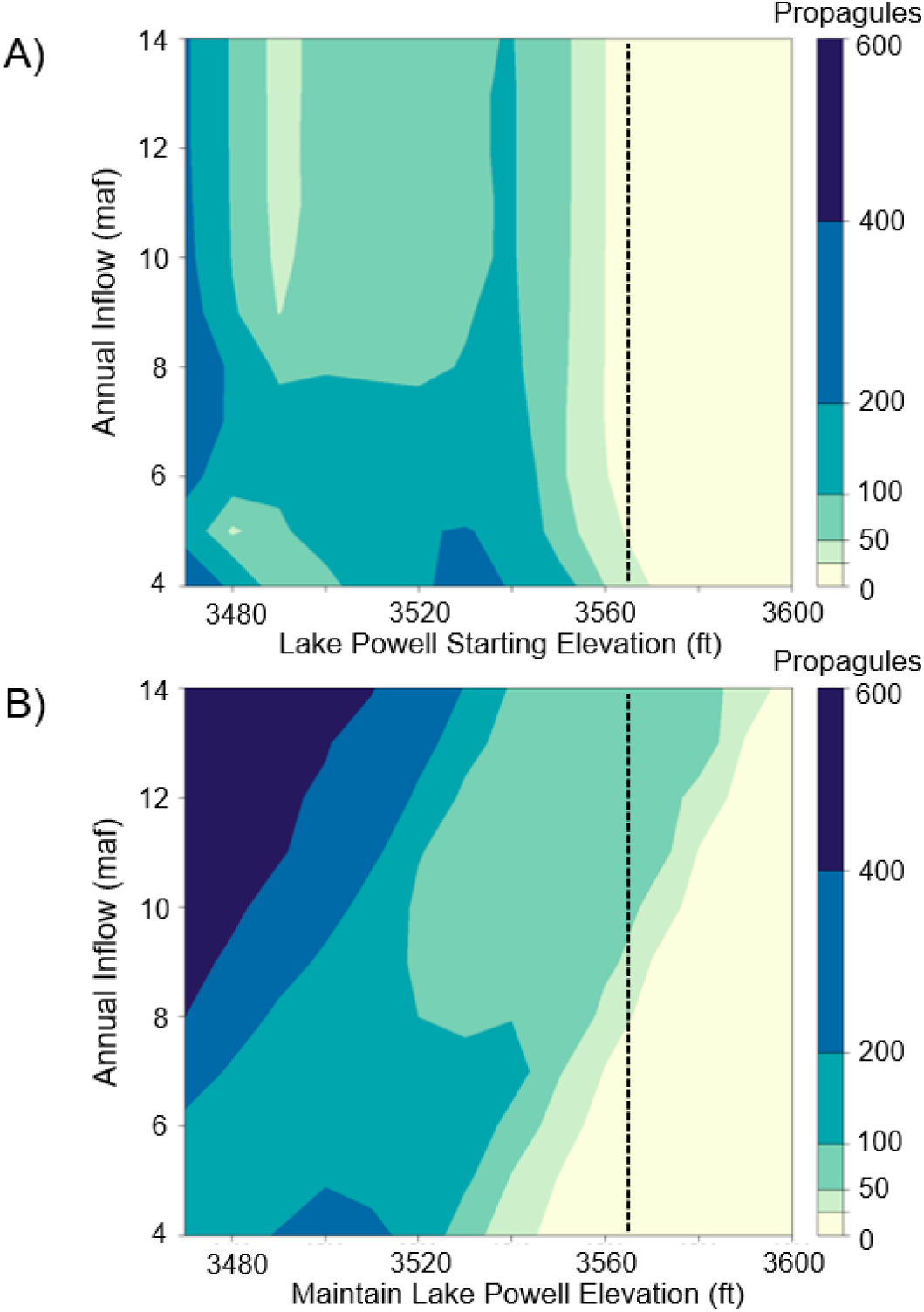
Estimated number of smallmouth bass entrained under different conditions. Predicted mean annual propagule pressure (indicated by shading in color ramp) from Lake Powell into the Lees Ferry tailwater based on Lake Powell elevations in feet (x axis) and annual inflow volume in million-acre feet (y axis) using A) predicted outflows for 2024, or B) maintaining elevation with outflows matching inflows at a monthly scale. Vertical, black dashed line denotes 2024 water year starting elevation.

**Figure 3:**
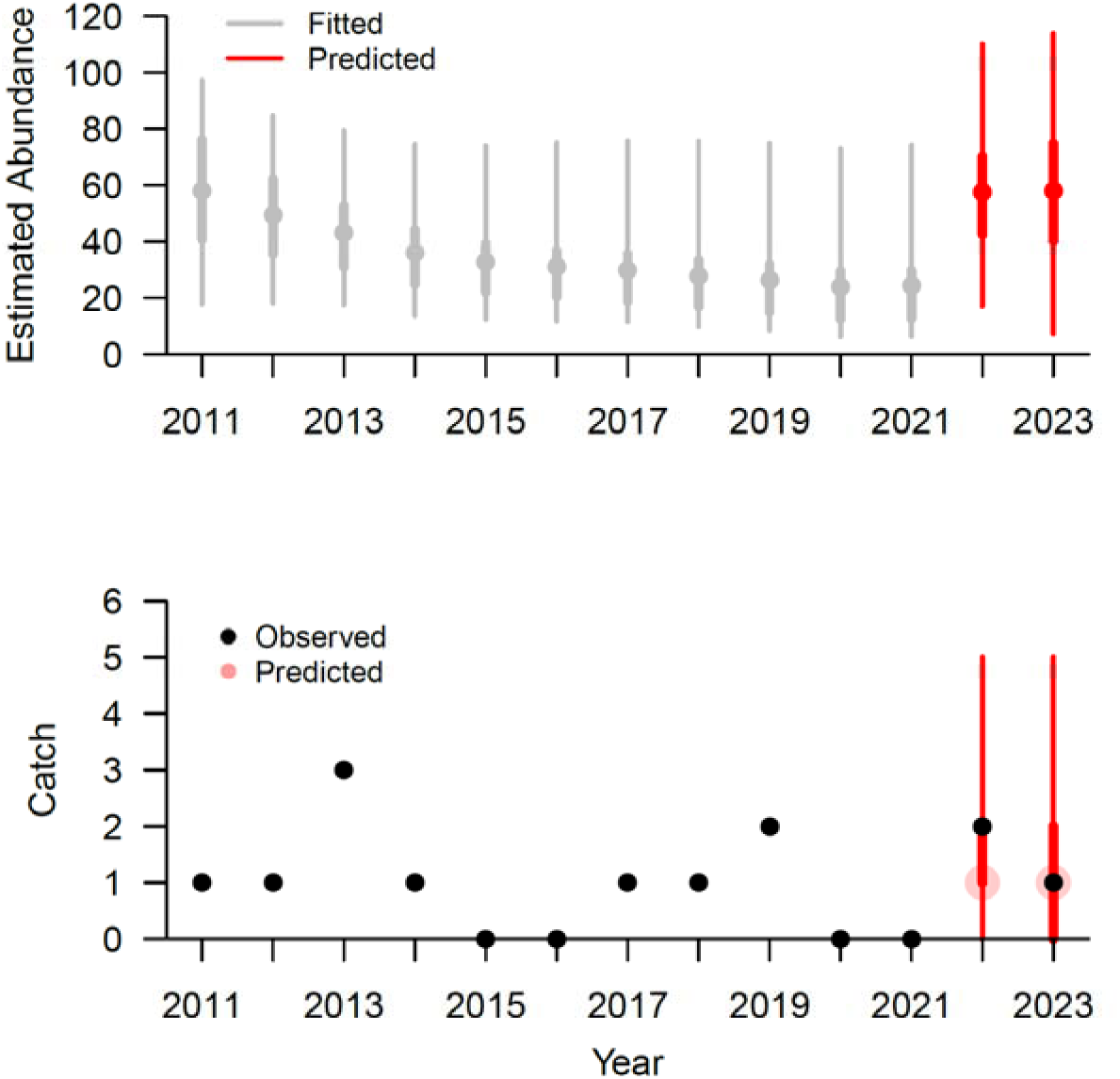
Estimates of the adult population size of smallmouth bass in Lees Ferry derived from the entrainment model fit to data from 2011 to 2021 and forecasted into 2022 and 2023 suggest a modest increase in adult smallmouth bass abundance during 2022 coincident with historically low reservoir elevations. Observed catch of adult smallmouth bass in 2022 and 2023 were consistent with these forecasts. For values that are estimated or predicted light whiskers cover the 95% credible intervals, while thicker whiskers cover the 50% credible intervals and dots indicate the median expectation. Black dots are observed data.

Release temperature from Glen Canyon Dam is primarily driven by reservoir elevation, and hence, the depth of withdrawal, time of year, and inflow volume (Figs. 4; S1). Temperatures downriver are dictated by release temperature, release volume, and solar insolation (Mihalevich et al., 2020), which drive increased thermal suitability for smallmouth bass farther from the dam (Dibble et al., 2021) (Figs. 4A; B; C). Based on the relatively moderate 2024 starting elevation (3,565 ft; 1,087 m), if all water is released through the penstocks over the course of the year, thermal conditions are unlikely to be suitable for population growth in the Lees Ferry tailwater but are likely to be suitable at the Little Colorado River confluence and highly likely to be suitable at the Diamond Creek confluence (Fig. 4A). Over a range of inflows, our model predicts that thermal conditions can be suitable for smallmouth bass population growth when Lake Powell starting elevations are less than 3,580 ft (1,091 m) and 3,600 ft (1,097 m) for Lees Ferry and the Little Colorado River confluence, respectively (Fig. 4B). However, the Diamond Creek confluence reach was thermally suitable under all elevations and outflow scenarios we evaluated (Fig. 4B). While large inflows increase elevations, they can also alter thermal profiles in Lake Powell resulting in greater variability of discharge temperature, including increasing the depth at which warm water occurs (Fig. S1), which can increase smallmouth population growth rates even at relatively high reservoir levels (Figs. 4B, C). This is most notable in our scenario that maintains constant Lake Powell elevation (Fig. 4C), where large annual inflows can create thermally favorable conditions for smallmouth bass even at relatively high reservoir elevations by pushing warmer surface water to deeper depths. However, when only considering smaller annual inflows (<8 million acre-feet: 9,868 million cubic meters), maintaining lower elevations at constant levels can likely result in no population growth at the Lees Ferry tailwater reach (Fig. 4C). Over the last fifty years, the only years for which our model predicts temperatures suitable for smallmouth bass reproduction were 2022 and 2023. Although monitoring efforts are not directly comparable across all of these decades, the tailwater has been well monitored for roughly thirty years (Eppehimer et al., 2024), and these are the only two years in which reproduction was observed.

**Figure 4:**
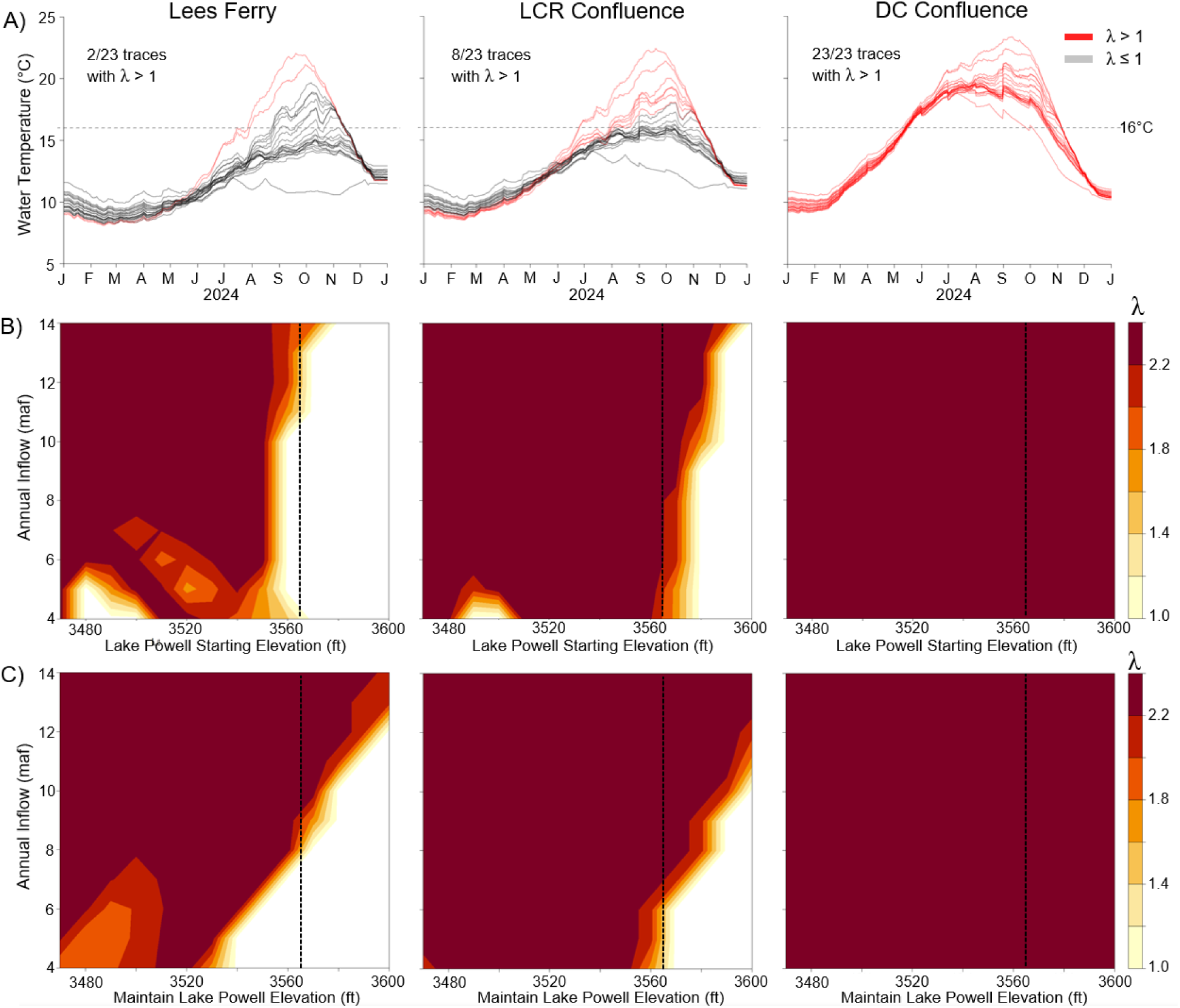
Thermal regimes and smallmouth bass population growth (λ) potential under different reservoir conditions, reservoir releases volumes, and geographic locations. A) Predicted daily water temperature for 2024 at Lees Ferry, Little Colorado River (LCR) confluence, and Diamond Creek (DC) confluence locations based on Lake Powell elevation projections using 23 different historic inflows. Red and grey lines indicate traces with and without predicted smallmouth bass population growth (λ), respectively. Horizontal dashed line denotes 16°C estimated smallmouth bass spawning threshold. Predicted λ (population growth rate) at each location based on Lake Powell elevations in feet (x axis) and annual inflow volume in million-acre feet (y axis) using B) predicted outflows for 2024, or C) maintaining elevation with outflows matching inflows at a monthly scale. Vertical, black dashed line denotes 2024 water year starting elevation. Note: Some traces drop below 3,490 ft (1,064 m) elevation triggering releases from the deeper river outlets, which reduces water temperature and λ.

## Discussion

Our models indicate that under current elevations and operations, the risk of smallmouth bass establishment in the Colorado River downstream from Glen Canyon Dam is high because low reservoir elevations increase both propagule pressure and thermal suitability of the river for reproduction and recruitment. Our results also suggest that temperatures will likely be favorable for smallmouth bass population growth near the Little Colorado River, and in western Grand Canyon near Diamond Creek, the two epicenters of threatened humpback chub populations in the lower basin. Based on long-term climatic trends of decreasing annual precipitation, Lake Powell elevations are likely to remain in the range we analyzed unless consumptive water use is reduced to match water supply in the Colorado River basin (Wheeler et al., 2022). Future runoff is expected to remain low, and water managers have struggled to reach consensus on how to balance supply and demand (Wheeler et al., 2022). Unless this imbalance is resolved, reservoirs like Lake Powell and Lake Mead are likely to maintain low water-storage conditions that threaten water security and native fish communities through non-native invasions. In the short-term, prioritizing water storage in Lake Powell and/or releasing water from the deeper river outlets may help minimize the risk of smallmouth bass establishment in the Grand Canyon segment of the Colorado River. Managers are currently exploring mid-term (3+ years) changes to infrastructure that could maintain cool temperatures and low rates of entrainment under lower reservoir elevation (US Bureau of Reclamation, 2023a; 2023b), however, over longer time periods, the water supply-demand imbalance that has drained basin reservoirs will likely need to be addressed (Schmidt et al., 2023) to avoid conditions that facilitate smallmouth bass invasion and establishment.

Our modeling suggests that if reservoir elevations are less than 3,540 ft (1,076 m) for significant portions of the year (Fig. S2), controlling the invasion of smallmouth bass will be challenging because entrainment of reservoir fishes will remain high so long as water is passed through the penstocks due to greater overlap with higher layers of the water column favored by smallmouth bass. Our model predicts that keeping reservoir starting elevations above 3,565 ft (1,087 m, 35% of capacity) would significantly reduce the risk of entrainment of additional smallmouth bass and other undesirable non-native species under current dam operations. If Lake Powell was managed to maintain a constant elevation with inflows matching outflows and annual inflow volumes were low, then elevations below that 3,565 ft (1,087 m) threshold could limit smallmouth bass propagule pressure.

Our models also indicate that under current dam operations keeping reservoir starting elevations above 3,600 ft (1,097 m, 47% of capacity) would likely create thermal conditions unsuitable for smallmouth bass from Glen Canyon Dam to the confluence with the Little Colorado River. Stopping the expansion of smallmouth bass is important for managers as the Little Colorado River provides critical habitat for humpback chub (Yackulic et al., 2014), and downstream warming limits the impact of Lake Powell elevation management in sections of western Grand Canyon. If future inflows are low, lower elevations could prevent population growth if reservoir releases were timed to maintain maximum elevation throughout the year by matching outflows with inflows. Given present and predicted water supply, higher elevations could likely only be achieved by prioritizing storage in Lake Powell over Lake Mead and lowering basin-wide demand (Wheeler et al., 2022). If reservoir storage in Lake Powell cannot be maintained above these critical levels and preventing smallmouth bass is an identified management priority for reservoir operations, during certain seasons when water releases are suitable for smallmouth bass spawning, water may need to be released through the deeper river outlets to cool the river in an effort to limit reproduction and recruitment.

Although creation of unsuitable thermal regimes could theoretically limit smallmouth bass population growth, eliminating invasion risk will also require limiting entrainment and downriver dispersal. Although our model suggests entrainment was modest in 2022 and 2023, it predicts substantially higher rates at elevations below 3,520 ft (1,073 m; Fig. 2). Over the past two decades prior to 2022, systematic sampling in Lees Ferry only captured 12 smallmouth bass; 10 of which were adults (>200 mm) (Eppehimer et al., 2024). However, in 2022, 69 age-0 smallmouth bass were captured in routine monitoring in that same reach in September, and an additional 269 age-0 fish were captured in targeted efforts in Lees Ferry from July to December (unpublished data: L. Tennant, NPS). The small size of some captured individuals (some 20 mm total length) suggests in situ reproduction occurred in Lees Ferry, which has not been documented previously, and was consistent with model forecasts of thermal conditions suitable for reproduction and population growth. In 2022, no smallmouth bass were captured below the Lees Ferry reach. At the same time, captures of adult smallmouth bass did not increase appreciably during routine monitoring, which supports our model predictions of only modest entrainment in 2022 (see Supplemental Materials). If reproduced (or entrained) smallmouth bass disperse to and become established in the Western portions of the Grand Canyon near Diamond Creek, modification of Lake Powell release temperatures (i.e., use of river outlets) will likely not affect population growth because of the extent of warming that occurs downriver (Dibble et al., 2021).

Starting in 2022, management agencies began removal efforts targeting smallmouth bass in the Lees Ferry tailwater. Although overall risk of dispersal may be reduced by mechanical removal, it will not eliminate the risk, and based on efforts in the upper basin mechanical removal is part of a portfolio of management options required to try to control smallmouth bass populations (Breton et al., 2014). Mechanical removal efforts are more likely to reverse the early stages of invasion if combined with cooler water temperatures that limit further reproduction and measures that limit further entrainment. Once a non-native fish species has become established, the benefits of mechanical removal are often short-lived, transitory, and at times non-existent, even when done at regular intervals and/or with high initial reduction estimates (Zelasko et al., 2016; Lennox et al., 2018; Michel et al., 2020). This is especially true for more open systems, which often allow a source of replenishment for nonnatives (Zelasko et al., 2016). In addition, mechanical removal efforts are often costly and have high logistical demands for management agencies (Runge et al., 2018). However, mechanical removal efforts have been used previously for rainbow trout in the Colorado River near the Little Colorado River (Coggins et al., 2011), as well as for brown trout in Bright Angle Creek (Healy et al., 2020) with variable success (Coggins et al., 2011).

Our entrainment model predicts overlap between the dam withdrawal zone in the reservoir and water column layers occupied by fish leading to greater rates of fish passage. This is supported by previous work suggesting successful fish passage is primarily determined by fish behavior and reservoir operations (Harrison et al., 2019). In some contexts, dam entrainment can be high enough to potentially impact fish populations both up and downstream (Harrison et al., 2019) and can vary significantly by species, life stage, and season (Martins et al., 2013; Lin et al., 2022); fine-tuning entrainment model parameters to describe specific contexts may be most informative. Additional research quantifying seasonal use of reservoir depths in the forebay of Glen Canyon Dam (underway), quantifying life stage-specific survival rates during entrainment, and quantifying outflow specific entrainment rates could be incorporated into the model structure to yield more precise forecasts.

Now that smallmouth bass are present in significant abundances downstream of Glen Canyon Dam, the success of their invasion will likely be driven by a multitude of abiotic factors (Moyle and Light, 1996) with river temperatures largely determining their potential for long-term establishment and population growth, as these temperatures govern reproduction. Temperature is a known driver of smallmouth bass population dynamics in the upper Colorado River basin (Breton et al., 2015; Bestgen and Hill, 2016) and throughout North America (Shuter et al., 1980). Ability to spawn and timing of spawn, as well as overwinter survival of age-0 fish are primarily tied to temperature (Shuter et al., 1980), and therefore temperature will likely determine the long-term success of an invasion. For example, in the John Day River in the Pacific Northwest USA, cooler water temperatures limited the upstream expansion of smallmouth bass (Rubenson and Olden, 2017). Warmer temperatures also benefit the native fishes of the Colorado River. Recent increases in humpback chub abundance in Western Grand Canyon (Yackulic et al., 2014; Van Haverbeke et al., 2017) are likely a result of warming river temperatures which substantially increase humpback chub growth rates (Dzul et al., 2017; Yackulic et al., 2018). However, any of these benefits from warmer water could be jeopardized by warm water driven smallmouth bass expansion (Dibble et al., 2021). This creates a complicated interaction between the two species where smallmouth bass could be predators on juvenile humpback chub, but warming water could expand humpback chub spawning and rearing habitat and would lead to faster growth rates, therefore reducing the vulnerability window for smallmouth bass predation.

There are other factors in addition to temperature that can limit smallmouth bass populations. Turbidity is highly variable spatially and temporally in the Colorado River between Lees Ferry and Diamond Creek, often depending on tributary inputs including the Paria and Little Colorado rivers (Topping et al., 2000). Turbidity has been shown to affect the incidence of pisciviory in rainbow and brown trout (Yard et al., 2011) and may influence predation vulnerability for humpback chub (Ward et al., 2016). Smallmouth bass are visual predators that forage more efficiently in clear water (Brown et al., 2009), and high turbidity and total suspended solids can decrease foraging efficiency and can impact survival, especially at early life stages (Sweka and Hartman, 2003; Suedel et al., 2017). The upper Colorado River is also a highly turbid system, turbidity regimes are similar between portions of the upper Basin that have been invaded and the Colorado River below Lees Ferry, and smallmouth bass have successfully invaded the upper Basin. Smallmouth bass nesting and early life survival can also be impacted by flow fluctuations (Winemiller and Taylor, 1982; Lukas and Orth, 1995; Knotek and Orth, 1998) and in the upper Colorado River basin flow fluctuations are currently being assessed as a management option to reduce smallmouth bass recruitment (Bestgen, 2018). Flow fluctuations could be a management option for reducing smallmouth bass recruitment between Lees Ferry and Diamond Creek.

Managing Lake Powell elevations at higher levels (>3,570 ft; 1,088 m) would likely minimize propagule pressure from the reservoir and would likely create downstream conditions that reduce the risk of smallmouth bass reproduction in the tailwater reach. However, higher elevations will likely be needed to create thermally unsuitable conditions farther downstream. Lambdas for smallmouth bass are potentially high and likely due to in situ reproduction rather than passage through the dam; prior to 2022 relatively few adult fish were caught in the Lees Ferry reach and were likely the result of entrainment, whereas fish caught beginning in 2022 were mostly sub-adults that were likely spawned locally. The periodic presence of turbidity in the lower reaches from tributary inputs may reduce smallmouth bass recruitment, but the Lees Ferry tailwater reach will always have relatively low turbidity. Smallmouth bass are not yet well established downstream of Glen Canyon Dam, but suitable temperatures, flow, habitat, and periodic low turbidity are likely to provide suitable conditions for spawning that is likely to result in future population increases.

Water supply and storage decisions directly impact river suitability for fishes, and certain reservoir operations may spread non-native fishes. Current water shortages throughout the Colorado River basin and corresponding unprecedented low reservoir levels in Lake Powell increase the risk of smallmouth bass establishment in Grand Canyon, which threatens native fishes. Forecasting biological invasions is crucial for management efforts, however, such forecasts are often unavailable. Predictive tools, such as the models presented here, can be useful for informing time-sensitive management decisions involving long-term water storage strategies as well as efforts to control high-risk non-natives over shorter time periods. For example, these models are currently being used to inform water storage decisions and designer flow releases in the Colorado River basin (US Bureau of Reclamation, 2023a; 2023b). Although specific to smallmouth bass and Glen Canyon Dam, our models provide a template that can be adapted to other species in different systems and can help managers address the novel challenges presented by long-term drought in regulated river systems.

## Acknowledgements

We received funding from the U.S. Geological Survey (USGS) Southwest Climate Adaptation Science Center, the USGS Ecosystem and Water mission areas, and the Bureau of Reclamation. We thank numerous managers who provided their input during early iterations of our model development. We also thank D. Ward, P. Budy, B. Friesen, A. Schultz, M. Anderson, W. Pine, R. Valdez, G. Chong, and M. Rumpf for their helpful reviews of this paper. The findings and conclusions in this article are those of the authors and do not necessarily represent the views of the U.S. Fish and Wildlife Service. Any use of trade, firm, or product names is for descriptive purposes only and does not imply endorsement by the U.S. Government.

