## Supplemental Materials for "Declining reservoir elevations following a two-decade drought increase water temperatures and non-native fish passage facilitating a downstream invasion"

**Supplemental Material**

Additional information and details on the models including parameterization are presented here and associated data and code are available from the USGS data release (Eppehimer et al., 2024).

*Forecasting reservoir elevations*

Both our entrainment and population growth models depend on forecasts of reservoir elevation. Forecasts of Lake Powell reservoir elevation (storage) are calculated at a monthly time step and based on elevation (storage) at the end of the preceding month, inflows, outflows, bank storage, and evaporative losses. The Bureau of Reclamation provides these projections in their ‘24-month study’ (available at: <https://www.usbr.gov/lc/region/g4000/riverops/24ms-projections.html>) using the Colorado River Simulation System (CRSS: Schuster, 1998; Wheeler et al., 2019), and when appropriate we used these projections. However, to allow us to consider Lake Powell inflows and outflows beyond the scenarios considered in the ‘24-month study,’ including future scenarios, we rewrote the code for the Lake Powell node in CRSS in R version 4.1.2 (R Core Team, 2021) allowing us additional flexibility. Lake Powell elevation to volume relationships were calculated at 0.1 ft (0.03 m) increments. We compared projected elevations from the ‘24-month study’ to output from our code using the same assumptions to confirm that our elevation projections were consistent.

*Lake Powell thermal profiles*

To predict the expected water temperature of releases, *T_d,z_*, on day, *d*, of the year when water is being drawn from depth below water surface, *z*, we fit the following model at 10-ft (3-m) depth intervals from 1 ft to 300 ft (0.3-91 m) deep:

Equation S1:	$T_{d,z}=\alpha_{d,z}{+\beta}_{d,z}I$

Equation S2*:*	$\alpha_{d,z}\sim N(\alpha_{d-1,z},\sigma_{\alpha,z})$

Equation S3:	$\beta_{d,z}\sim N(\beta_{d-1,z},\sigma_{\beta,z})$

where *I* is the standardized inflow in the prior year when *d* is earlier than June 1^st^ and in the current year when *d* is June 1^st^ or later, *α_1,z_* was given a U(5,25) prior and *β_1,z_* was given a U(-5,5) prior. We fit this model to data extracted for a given depth from 225 temperature profiles collected between 2000 and 2022 (Andrews and Deemer, 2022). Fitting occurred in *Stan* (Stan Development Team, 2022a) using three chains, each with 1,000 sampling iterations. We then linearly interpolated $\alpha_{d,z}$ and $\beta_{d,z}$ values for each foot bin between the 10-ft (3-m) depth models allowing us to predict water temperature for each day of the year at a 1-ft (0.3-m) resolution. We assumed that temperatures at depths >300 ft (91 m) were the same as 300 ft (91 m). To predict release temperatures on day, *d*, of a given year, we chose the value in $T_{d,z}$ given by that day and the corresponding depth, *z*, consistent with the centerline of either the hydropower or bypass tubes given the forecasted reservoir elevation on that day (Fig. 1B).

*Entrainment model details*

To quantify the risk of smallmouth bass establishment based on propagule pressure in the river downstream of Lake Powell, we developed a model that coupled abundances in Lake Powell, $N_{y, d}^{Res}$, in year, *y*, on day, *d*, to abundances in the river, $N_{y}^{Riv}$ via the following equations:

Equation S4:	$N_{y,d+1}^{Res}=N_{y,d}^{Res}*(1-\gamma(E_{y,d}-E_{out}))$

Equation S5:	$N_{y+1}^{Riv}=\varphi_{ann}*N_{y}^{Riv}+\sum_{d=1}^{365} N_{y,d}^{Res}*\gamma\left( E_{y,d} \right)*\varphi_{ent}-M_{y+1}$

Where $\gamma$ is a function (described in more detail in the following paragraph) that estimates how the entrainment rate, on a per fish basis, varies with the depth of the outflow structure in the reservoir (calculated as the difference between the reservoir elevation, $E_{y,d}$, and the elevation of the center of the outflow structure, $E_{out}$), $\varphi_{ann}$ is the survival rate of smallmouth already present in the river (from entrainment in prior years), $\varphi_{ent}$ is the survival rate of smallmouth passing through Glen Canyon Dam, and $M_{y+1}$ is the number of smallmouth bass captured in the river (and removed) – for all years used in fitting the model these fish were captured through regular monitoring; however, beginning in 2022 targeted removal activities occurred in addition to regular monitoring. In fitting the model, we used counts of $M_{y+1}$ from the past 10 years (2011-2021) since monitoring has been relatively constant during this period and each year’s number of smallmouth bass captured and removed in Lees Ferry has varied between 0 and 3 (Eppehimer et al., 2024). Specifically, we linked our models of abundance to observed counts via the equation $M_{y+1}\sim Pois({p*N}_{y+1}^{Riv})$, where $p$ was given a uniform prior between 0.005 and 0.05.

The exact form of the function, $\gamma\left( d \right)$, describing entrainment as a function of the depth of the centerline of the outflow structure, *d*, is given by:

Equation S6:	$\gamma\left( d \right)= \sum_{d-12}^{d+12} e^{\mu_{\gamma}+\alpha_{\gamma}*\rho_{d}}$

Where $e^{\mu_{\gamma}+\alpha_{\gamma}*\rho_{d}}$ represents the probability that smallmouth bass are distributed within depth bin *d* and summing occurs from 12 ft (3.7 m) above to 12 ft (3.7 m) below the centerline of the outline to represent the diameter of the intake. $\alpha_{\gamma}$ and $\mu_{\gamma}$ are estimated parameters while $\rho_{d}$ represents the expected temperature suitability of each depth from 1 ft to 301 ft (0.3-91 m) below the surface with 1-ft (0.3 m) resolution calculated from 225 temperature profiles collected between 2000 and 2022 (Andrews and Deemer, 2022), via the following equation:

Equation S7:	$\rho_{i}=\frac{\sum_{y=1}^{225} \Phi(T_{i,y})}{\sum_{j=1}^{301} \sum_{y=1}^{225} \Phi(T_{j,y})}$

Where $T_{i,y}$ represents the temperature at depth *i* on profile *y* and $\Phi$ is a function that returns a zero when $T_{i,y}\leq16$, a value of 1 when $T_{i,y}\geq22$, and a value of ($T_{i,y}-16)/6$ when $16<T_{i,y}<22$ derived from temperature suitability models for smallmouth bass described by Dibble et al. (2021). To improve estimates of $\mu_{\gamma}$ and $\alpha_{\gamma}$ we extracted the number of detections of fish in 13 3-4 m depth bins during hydroacoustic surveys conducted between 2007 and 2009 near the forebay of Glen Canyon Dam and reported in Figure 6 from Bureau of Reclamation (2012). We assumed that the sum of $e^{\mu_{\gamma}+\alpha_{\gamma}*\rho_{d}}$ within each depth bin divided by the overall sum of $e^{\mu_{\gamma}+\alpha_{\gamma}*\rho_{d}}$ across all depths would predict the multinomial distribution of detections across depth bins. Briefly, this Bureau of Reclamation report found that 69% of fish occupied the top 4 m with 98% of fish found within 14 m of the surface, and we assumed smallmouth bass followed these same distributions. We acknowledge that these surveys did not differentiate species. However, similar results in the literature suggest that smallmouth bass often occupy the top portion of the water column (e.g., 1-5 m depth) but can go deeper when temperatures are not stratified (e.g., 14 m depth; Suski & Ridgway, 2009).

We developed a prior for $N_{y, d=1}^{Res}$, the size of the Lake Powell smallmouth bass population size that is proximate to the dam, and thus capable of being entrained from the following assumptions regarding smallmouth bass densities and home range areas derived from analysis of published literature. In North America, smallmouth bass reservoir densities range from 0.74 to 164 per hectare (ha; 0.0074-1.64 km^2^) (Carlander, 1977), and home range areas range from 0.01 ha (0.0001 km^2^) (Savitz et al., 1993) to 302 ha (3.02 km^2^) (Ridgway & Shuter, 1996) with an average lower bound of 10.8 ha (0.108 km^2^), an average upper bound of 41.1 ha (0.411 km^2^), and an average mean of 22.2 ha (0.222 km^2^) (Savitz et al., 1993; Minns, 1995; Ridgway & Shuter, 1996; 2002; Brown et al., 2009; Smith et al., 2022). Given the relative lack of littoral habitat in Lake Powell (Pennock & Gido, 2021), we assumed smallmouth bass home ranges could extend along the canyon wall bound shoreline, thus estimates of movement could reach large distances (e.g., >10 km). Based on combinations of low/high densities and potential movement within small/large home ranges, we estimated that the smallmouth bass population coming into the dam forebay ranged from 1,000 to 50,000 individuals. At the beginning of each new year (*i.e.,* *d*=1), $N^{Res}$ is assumed to be reset to its initial value based on local recruitment and immigration from elsewhere in Lake Powell.

A prior for smallmouth bass entrainment survival, $\varphi_{ent}$, was estimated from a meta-analysis by Algera et al. (2020). Survival rates vary in relation to a multitude of factors including turbine design (Algera et al., 2020). Based on this available literature on fish passage (excluding eels) through Francis turbines (used in Glen Canyon Dam), mean survival was 0.588 (0.255 SD). This estimate from Algera et al. (2020) is an average of survival rates from 13 studies encompassing 14 species, of which >90% were salmonids. When reported, the average total length of studied fish was 138 mm, 61 mm SD. Our entrainment survival estimate could be biased low given that smallmouth bass are a relatively hearty fish (Brown et al., 2009) but could also be biased high given that high pressure head dams like Glen Canyon Dam were not included in analyses by Algera et al. (2020).

The annual survival rate of adult smallmouth bass, $\varphi_{ann}$, was given a prior of 0.73 with a standard deviation of 0.1. The mean value was derived from models describing smallmouth bass population dynamics in the Yampa and Green Rivers (Breton et al., 2015; Bestgen and Hill, 2016; see *Population growth rate model details*).

Our propagule pressure model assumes no seasonal variation in smallmouth bass with proximity to Glen Canyon Dam since no data were available when we were developing the model; however, alternate models could relax this assumption. In addition, the model does not account for local reproduction below the dam as local reproduction was not observed until 2022: all but two smallmouth bass observed in Lees Ferry prior to 2022 were adults (> 200 mm) (Eppehimer et al., 2024).

*Population growth rate model details*

Lake Powell inflows were based on resampling the past 23 years of data (2000-2022; available at: <https://www.usbr.gov/rsvrWater/HistoricalApp.html>). Lake Powell vertical thermal profiles (Andrews and Deemer, 2022; see *Lake Powell thermal profiles*) were related to forecasted reservoir elevation to predict the temperature of water released from Glen Canyon Dam. Our downriver temperature model adapted Dibble et al. (2021) to estimate daily rather than monthly average water temperature at different locations downstream of Glen Canyon Dam by using loess smoothed (span = 0.2) daily average values for air temperature (National Climate Data Center: USW00003162) and solar insolation (global horizon irradiance; National Solar Radiation Database: 97653) from national weather stations located in Page and Williams, AZ, respectively.

Survival and fecundity parameters used to forecast smallmouth bass values in response to variable river temperatures were derived from a smallmouth bass age-structured demographic model developed for the Green River sub-basin of the Colorado River (Breton et al., 2015; Bestgen and Hill, 2016; unpublished data available from: L. Bruckerhoff, OSU,). Age specific survival and fecundity were assumed constant for all ages except age-0 fish. Model structure was adapted from Breton et al. (2015) for the Yampa River and was expanded to include dynamics in the middle Green River reach between the confluences with the Yampa and White Rivers using long-term catch-effort, age, and growth observations (Bestgen and Hill, 2016; unpublished data available from: L. Bruckerhoff, OSU,). The model structure included movement between the Yampa and Green Rivers, reproduction, exploitation due to mechanical removal efforts, and annual survival. Parameters describing adult and subadult age specific survival, age-0 survival, fecundity, and capture probabilities were estimated by fitting our model to observed catch effort data for juveniles (age-0), sub-adults (ages 1-2) and adult (ages 3+) smallmouth bass from the Lily Park and Yampa Canyon reach of the Yampa River and the Echo-Split reach of the Green River (2004-2021) (unpublished data available from: L. Bruckerhoff, OSU,). Catch-effort and abundance estimates were extracted from annual reports publicly available on the Upper Colorado River Endangered Fish Recovery Program website (<https://coloradoriverrecovery.org/uc/documents/work-plan-documents/annual-reports/>). Detailed descriptions of sampling methods can be found in these reports. Models were fit using Bayesian inference using MCMC sampling using Stan software version 2.31 (Stan Development Team, 2022a) implemented through R version 4.1.2 (R Core Team, 2021) using the package RStan version 2.21.7 (Stan Development Team, 2022b).

Sub-adult and adult survival were age specific based on a general size-mortality relationship described by Lorenzen (1996):

Equation S8:	*M_W_ = M_u_W^b^*

where *M_W_* is the natural mortality rate at weight (*W*), *M_u_* is the mortality rate per unit weight, and *b* is an allometric scaling factor. We estimated *M_u_* and *b* for each age class using mean observed lengths of smallmouth bass in each age class and estimated weights using the standard weight equation for smallmouth bass (Kolander et al., 1993). We constrained estimates of *M_u_* and *b* using informative priors derived from values reported in Lorenzen (1996).

Survival of age-0 fish was dependent on estimated end-of-summer length (based on estimated hatch dates and water temperatures post hatch) and winter water temperatures. We assumed a water temperature spawning threshold of 16°C daily average. We conducted a literature review of water temperatures during the initiation of smallmouth bass spawning. We refined our search by excluding studies conducted in hatcheries and excluding studies that lacked precise temperature records. Also, we only included studies that recorded observations of smallmouth bass eggs in nests (as opposed to pre-spawn behavior alone like nest building). If there were multiple studies on the same site, we chose the most recent. And if a study included multiple sites or multiple years with different temperature thresholds, we chose the lowest temperature reported. This review of the literature showed spawning initiation thresholds ranging from 15 to 16.2°C daily average water temperature (Table S1). Based on 2003-2011 data from the Yampa and Green Rivers of the upper Colorado River basin, Bestgen and Hill (2016) estimated a smallmouth bass spawning threshold of 16°C using otolith aging of age-0 fish. We chose 16°C as an estimated spawning threshold in our models because it fell within the range of our literature review, and the regulated and partially regulated river reaches studied by Bestgen and Hill (2016) most closely matched the Colorado River between Lake Powell and Lake Mead.

We predicted hatch dates based on regression models presented in Bestgen and Hill (2016) and based off of 20 years of age-0 data from the Yampa and Green Rivers (available at: <https://coloradoriverrecovery.org/uc/documents/work-plan-documents/annual-reports/>), where the date of first hatch is a function of the date water temperatures first reach 16°C daily average plus 7 days, during which time the average daily temperature does not drop below 13.9°C, which was the lowest observed spawning temperature in the upper Colorado River basin following onset of 16°C water temperatures. We then used age-0 growth regressions developed in Bestgen and Hill (2016) and modified by L. Bruckerhoff (unpublished data available from: L. Bruckerhoff, OSU,) to predict total length (TL) of age-0 fish going into winter based on the hatch date ($x_{t}^{hatch date}$) and river temperature ($x_{t}^{temperature}$ ) during the period of critical growth (45 days post hatch):

Equation S9: TL_t,i_ = 134.1 + ($x_{t}^{hatch date}$*-0.66) + ($x_{t}^{temperature}$* 2.82) + ($x_{i}^{day of hatch}$ * -1.6)

where $x_{i}^{day of hatch}$ is the day of hatch (1 through 31). Total lengths of age-0 fish were then used to predict overwinter survival (φ^age 0^_t_*)* as:

Equation S10:	φ^age 0^_t_ = β^0,winter^ + (β^1,winter^ *TL_,t_ ) + (β ^2,winter^ * s_days_,t_)

where *β^0,winter^* is the intercept for age-0 survival, *β^1,winter^* is the slope for the effect of total length of age-0 fish in each cohort (*TL_t_*), and *β ^2,winter^* is the slope for the number of starvation days during the winter (*s_days_t_*). The number of starvation days was calculated by counting the number of days mean daily river temperature was less than or equal to 10°C. Also, any fish with a predicted total length below 30 mm was assumed to not survive to the next year regardless of winter water temperatures. All slope and intercept parameters were estimated using the Green River sub-basin model and mean parameter estimates were used in the λ projections presented in the main text of this paper (β^0,winter^ = -2.14, β^1,winter^=2.66, β ^2,winter^ =-0.25).

Fecundity (defined here as the number of offspring produced per reproductive adult) was based on estimates of the intrinsic growth rate estimated from smallmouth bass data in the Green and Yampa Rivers (available at: <https://coloradoriverrecovery.org/uc/documents/work-plan-documents/annual-reports/>, unpublished data available from: L. Bruckerhoff, OSU,) and assumed smallmouth bass reproduction was not affected by densities in Lees Ferry. Specifically, we assumed an estimated fecundity of 51.15 offspring/adult with 63% of age 3 fish and 75% of age 4 and older fish reproducing in our λ projections (Breton et al., 2015), and we held this value constant across simulations.

Any use of trade, firm, or product names is for descriptive purposes only and does not imply endorsement by the U.S. Government.

**Supplemental Figures**

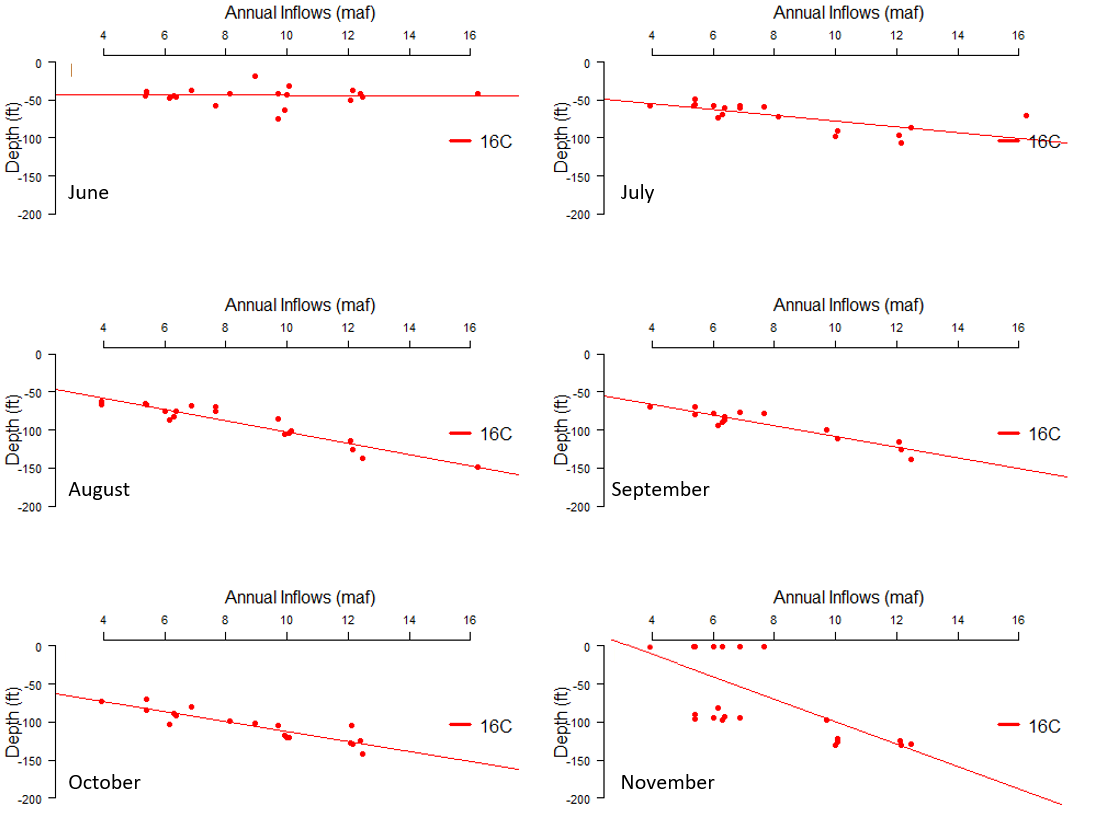

**Figure S1** Month specific Lake Powell thermal profiles showing the depth below surface (ft) at which water temperature is 16°C under different annual inflow scenarios (in million acre feet; maf) based on historic profiles from 1999-2020. Red dots show 16°C depth, and the red line is the linear line of best fit, which estimates the depth to which temperatures are ≥16°C. Plots illustrate these profiles for June through November.

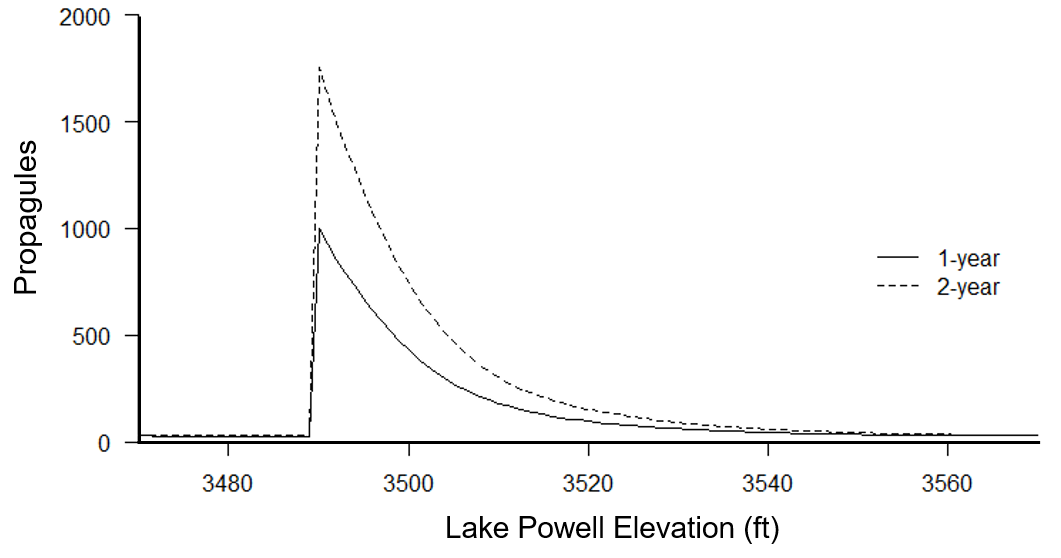

**Figure S2** Mean number of smallmouth bass propagules that persist over one- and two-year periods (solid and dashed lines, respectively) as predicted by the entrainment model (y axis) given constant Lake Powell elevations in ft (x axis). Increases from one- to two-year period are indicative of propagule pressure that surpasses adult mortality. At elevations at and below 3,490 ft (1,064 m), Glen Canyon Dam releases water from the deeper river outlets resulting in an abrupt decrease in predicted propagules depicted above.

**Supplemental Table**

**Table S1** Summary of temperature at which smallmouth bass initiate spawning from the literature including only direct observations of eggs in nests and excluding studies in hatcheries. For multiple studies on a single site, we chose the most recent, and for multiple sites within a single study, we chose the lowest temperature reported. The table includes daily average water temperature in °C, location of study, whether observed spawning was in river or lake, and citation.

| **Temperature (°C)** | **Location** | **Type** | **Citation** |
| --- | --- | --- | --- |
| 15 | Nagano, Japan | Lake | Peterson & Kitano, 2022 |
| 15 | Oregon, USA | River | Rubenson & Olden, 2019 |
| 15.2 | Ontario, CA | Lake | Turner & MacCrimmon, 1970 |
| 15.5 | Saskatchewan, CA | Lake | Rawson, 1938 |
| 16.2 | Oklahoma, USA | River | Dauwalter & Fisher, 2007 |
